# Soluble VE-cadherin disrupts endothelial barrier function via VE-PTP/RhoA signalling

**DOI:** 10.1101/2022.10.17.512494

**Authors:** Juna-Lisa Knop, Natalie Burkard, Mahshid Danesh, Thomas Dandekar, Mugdha Srivastava, Matthias Hiermaier, Jens Waschke, Sven Flemming, Nicolas Schlegel

**Affiliations:** University Hospital Wuerzburg, Department of General, Visceral, Transplantation, Vascular and Paediatric surgery (Department of Surgery I), Oberduerrbacherstraße 6, D-97080 Wuerzburg, Germany; Chair of Vegetative Anatomy, Institute of Anatomy, Faculty of Medicine, LMU Munich, Munich, Germany; University of Wuerzburg, Department of Bioinformatics, Biocenter, Am Hubland, D-97074 Wuerzburg, Germany; Core Unit Systems Medicine, 97080 Würzburg, Germany

**Author notes:** both authors contributed equally to this work (Juna-Lisa Knop, Natalie Burkard). both authors contributed equally to this work (Sven Flemming, Nicolas Schlegel). Corresponding author: Prof. Dr. Nicolas Schlegel, Department of Surgery I, Oberdürrbacherstraße 6, D-97080 Würzburg, Germany.

**Keywords:** endothelial barrier function, soluble VE-cadherin, inflammation, VE-PTP, endothelial junction

## Abstract

**Aim:** Increased levels of soluble Vascular endothelial (VE)-cadherin fragments (sVE-cadherin) have previously been linked with inflammation-induced loss of endothelial barrier function. We tested whether sVE-cadherin is critically involved in the onset of endothelial barrier dysfunction.

**Methods and Results:** Application of recombinant human sVE-cadherin (extracellular domains EC1-5) on human microvascular endothelial cells *in vitro* and in a rat model *in vivo* induced loss of endothelial barrier function and reduced microcirculatory flow. sVE-cadherin^EC1-5^ led to decreased localization of VE-cadherin at cell borders. Additionally, sVE-cadherin^EC1-5^ perturbed VE-protein tyrosine phosphatase (VE-PTP)/VE-cadherin interaction. VE-PTP inhibitor AKB9778 blunted all sVE-cadherin^EC1-5^-induced effects *in vitro* and *in vivo*. Downstream effects involve VE-PTP-dependent RhoA activation which was attenuated by AKB9778. Rho-kinase inhibitor Y27632 blocked sVE-cadherin^EC1-5^-induced loss of endothelial barrier function.

**Conclusion:** sVE-cadherin disrupts endothelial barrier function by dismantling the VE-cadherin complex at cell borders via VE-PTP-dependent RhoA activation. This uncovers a novel pathophysiological role of sVE-cadherin in the context of endothelial barrier dysfunction in inflammation.

## Introduction

Organ failure in sepsis and in systemic inflammation have been recognized as the critical factors leading to high mortality rates in patients (Bauer et al., 2020). This is reflected by the definition of sepsis as a life-threatening organ dysfunction caused by a dysregulated host response to infection (Singer et al., 2016). From a pathophysiological point of view, organ failure is primarily caused by loss of the microcirculation which results in metabolic dysfunction (Deutschman and Tracey, 2014; Lupu et al., 2020; Schick et al., 2012). In this context, one of the most critical steps is characterized by the breakdown of endothelial barrier function leading to extravasation of fluid and loss of microcirculatory flow (Deutschman and Tracey, 2014; Ince et al., 2016; Parikh, 2017).

Under resting conditions, endothelial cells line all blood vessels to provide a semipermeable and selectively regulated barrier (Radeva and Waschke, 2018; Spindler et al., 2010). Under inflammatory conditions, increased paracellular permeability is caused by disruption of tight junctions (TJ) and adherens junctions (AJ) connecting neighboring endothelial cells (Radeva and Waschke, 2018; Spindler et al., 2010). The crucial determinant of endothelial barrier integrity is the 120 kDa vascular endothelial (VE)-cadherin which is a transmembrane Ca^2+^-dependent adhesion protein. VE-cadherin is usually found as a dimer with an intra- and extracellular domain, the latter consists of 5 subdomains (EC1-5) (Dejana and Vestweber, 2013). It has been reported that the integrity of endothelial barrier function is strictly dependent on the interaction between VE-cadherin and VE-protein tyrosin phosphatase (VE-PTP) (Vestweber, 2021). VE-PTP is a transmembrane protein which interacts with VE-cadherin via its extracellular domain with the EC5 domain of VE-cadherin (Nawroth et al., 2002). Under conditions of inflammation, the VE-cadherin/VE-PTP interaction is impaired leading to downstream signaling which induces loss of VE-cadherin-mediated adhesion and consecutive breakdown of endothelial barrier function (Frye et al., 2015). Therefore, this mechanism has not only been attributed to play a critical role to maintain microvascular barrier function under resting conditions but also contributes to breakdown of endothelial barrier function in systemic inflammation and sepsis (Vestweber, 2021).

Additionally, it has been demonstrated that VE-cadherin is cleaved under inflammatory conditions leading to the release of soluble VE-cadherin (sVE-cadherin) fragments which predominately consist of the EC1-5 domains (Flemming et al., 2015; Schulz et al., 2008). VE-cadherin cleavage is caused by the activation of sheddases including the specific disintegrin and metalloproteinase ADAM10 (Schulz et al., 2008). Interestingly, clinical data show that increased levels of sVE-cadherin in the blood of septic patients are associated with negative outcome of these patients (Flemming et al., 2015; Yu et al., 2019; Yu et al., 2021; Zhang et al., 2010). Therefore, in a previous study we suggested sVE-cadherin as a potential clinical marker for diagnosing endothelial dysfunction in inflammation (Flemming et al., 2015). This is supported by several studies which correlated elevated sVE-cadherin levels with the onset of different inflammatory diseases (Chen et al., 2014; Ostrowski et al., 2017; Yu et al., 2021).

Besides that, *in vitro* data led to the idea that the release of sVE-cadherin alone may severely contribute to loss of endothelial barrier function independent from the presence of pro-inflammatory stimuli (Flemming et al., 2015). This may point to a novel and yet unidentified pathophysiological role of sVE-cadherin to mediate or aggravate endothelial barrier dysfunction following the onset of inflammation. However, currently there is no direct evidence for this hypothesis. The aim of the present study was to specifically address the contribution of sVE-cadherin to loss of endothelial barrier function.

## Results

### sVE-cadherin^EC1-5^ induced a dose-dependent loss of endothelial barrier integrity in vitro

To test our hypothesis of a causal role of sVE-cadherin to contribute to loss of endothelial barrier function we generated recombinant sVE-cadherin EC1-5. The human sequence of sVE-cadherin, which contains only the extracellular domain EC1-5, was cloned into the pcDNA-pDEST47 vector. The resulting plasmid pDEST47-sVE-cadherin^EC1-5^ was transfected into CHO cells (Figure 1A). The transfection resulted in the secretion of sVE-cadherin^EC1-5^ in cell culture supernatants. The presence of sVE-cadherin^EC1-5^ in CHO cells transfected with pDEST47-sVE-cadherin^EC1-5^ was verified by sequencing (suppl. Figure 1), by its recognition in an ELISA that specifically detects sVE-cadherin^EC1-5^ at levels ranging at 240.22 ± 18.22 ng/ml. Western Blot analyses revealed a protein at 90 kDa indicating the presence of the extracellular domain of VE-cadherin (sVE-cadherin^EC1-5^)(Flemming et al., 2015) in supernatants and cell lysates of CHO cells transfected with pDEST47-sVE-cadherin^EC1-5^ whereas no band was detected in untransfected CHO cells (Figure 1B).

**Figure 1:**
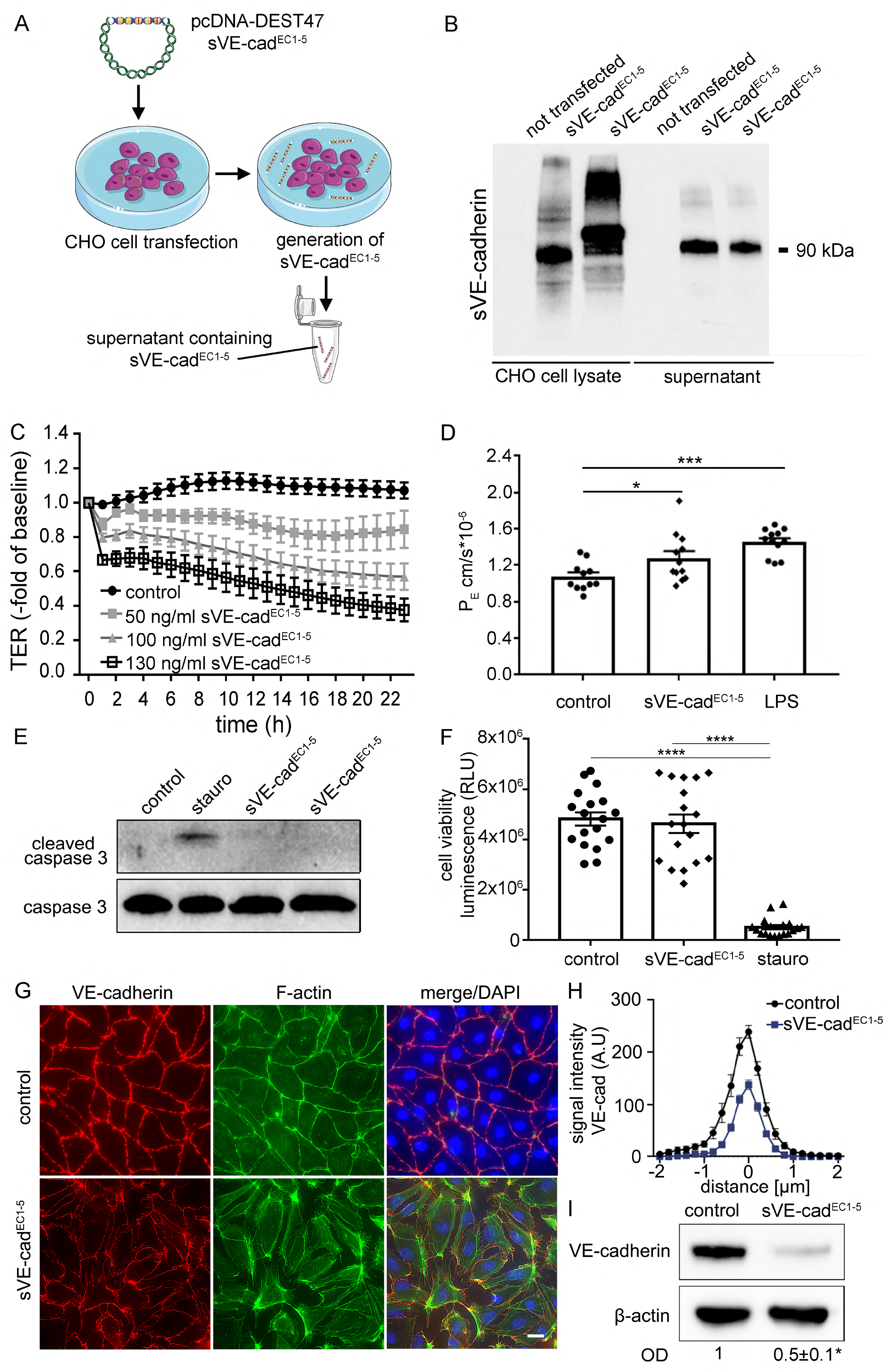
Generation and effects of sVE-cadherin ^EC1-5^ (sVE-cad^EC1-5^) on endothelial cells. A) Schematic overview of sVE-cad^EC1-5^ generation. CHO cells were transfected with the pcDNA-DEST47 plasmid containing sVE-cad^EC1-5^ and secrete sVE-cad^EC1-5^ into the supernatant. B) Representative Western Blot of CHO cells. In whole cell lysates of transfected CHO cells, a specific 90kDa protein band representing sVE-cadherin was detectable. Non-transfected CHO cells served as negative control. C) Application of sVE-cad^EC1-5^ on human endothelial cells (HDMEC) resulted in a dose-dependent decrease of transendothelial electrical resistance (TER). (p<0.0001, n=4 for each condition, one-Way ANOVA with Friedman and Dunn’s Test.) D) Permeability coefficient (PE) measured as 4kDa FITC dextran flux across confluent (HDMEC). sVE-cad^EC1-5^ resulted in a significant increase in P_E_ (p<0.038, n=3, one Way ANOVA with Sidak test). Lipopolysaccharide (LPS) served as a positive control (p<0.001, n=3, one Way ANOVA with Sidak Test). E) Representative Western Blot probed with cleaved caspase 3 antibody (used as marker of apoptosis.) Only HDMEC treated with staurosporine showed a specific protein band for cleaved caspase 3. In lysates of sVE-cad^EC1-5^ treated cells, no cleaved caspase 3 was detectable; n= 4 F) Cell viability assay showed that sVE-cad^EC1-5^ did not induce cell death (n=18, 1-way ANOVA). Staurosporine was used as positive control (p<0.0001, n=18, 1-way ANOVA). G) Representative immunofluorescence staining of HDMEC treated with sVE-cad^EC1-5^ for 24h. VE-cadherin staining in red) F-actin in green; experiment shown is representative for n>5. H) Quantifications of VE-cadherin immunostaining of all experiments for the conditions shown in G).Application of HDMEC with sVE-cad^EC1-5^ resulted in a significant redistribution of VE-cadherin from the cell border to the cytoplasm (p< 0.03). I) Representative Western Blot of HDMEC cell lysate probed with VE-cadherin antibody directed against the extracellular domain of VE-cadherin. Beta actin is shown as a loading control. The protein level is significantly reduced after application of sVE-cad^EC1-5^ (p<0.021, n=5, unpaired two-tailed t-test).

Next, we tested for effects of sVE-cadherin^EC1-5^ on endothelial barrier function using human endothelial cells (HDMECs). We used sVE-cadherin^EC1-5^ at different doses (50 ng/ml, 100 ng/ml and 130 ng/ml) and performed measurements of transendothelial resistance (TER) across confluent endothelial monolayers. Application of sVE-cadherin^EC1-5^ at 100 ng/ml and 130 ng/ml significantly reduced TER compared to controls to 0.57 ± 0.07-fold of control and 0.38 ± 0.06-fold of control after two hours whereas 50 ng/ml had no effect on endothelial barrier properties (Figure 1C). For the following experiments a dose of 130 ng/ml sVE-cadherin^EC1-5^ was chosen. In measurements of 70 kDa FITC-Dextran flux across endothelial monolayers following application of sVE-cadherin^EC1-5^ resulted in augmented permeability coefficients (P_E_) compared to controls after 1 h (1.10 ± 0.05 cm/s*10^-6^ for controls, 1.27 ± 0.08 cm/s*10^-6^ sVE-cad^EC1-5^). Comparable results were obtained after incubation of endothelial monolayers with 100 ng/ml LPS which increased P_E_ to 1.44 ± 0.06 cm/s*10^-6^ (Figure 1D). In cell viability assays we excluded that application of sVE-cadherin^EC1-5^ on endothelial cells resulted in the induction of cell death (Figure 1E, F). No induction of apoptosis was detected when endothelial cells were incubated with sVE-cadherin^EC1-5^ whereas staurosporine as a known inductor of apoptosis led increased cleaved caspase-3 (Figure 1E).

As revealed by immunostaining, VE-cadherin in confluent endothelial monolayers was regularly distributed at cell borders under control conditions whereas application of sVE-cadherin^EC1-5^ resulted in a fragmented staining pattern indicating loss of VE-cadherin (Figure 1G, H). This was paralleled by augmented stress fibers in F-actin staining and intercellular gap formation as well as by an overall reduction of VE-cadherin to 0.5 ± 0.1-fold of controls as revealed by Western blotting.

Next, we investigated proteins known to be associated with VE-cadherin such as α-, β-, and γ-catenin at the cell borders under control conditions. sVE-cadherin^EC1-5^ treatment of endothelial cells led to a reduction of all these proteins at cell borders (Figure 2A, C, E). Similarly, the TJ-associated protein ZO-1, which was also found to be regularly distributed at cell borders, was diminished following sVE-cadherin^EC1-5^ application (Figure 2G). The protein levels of AJ-associated proteins α-, β-, and γ-catenin and ZO-1 were unchanged following incubation of endothelial cell with sVE-cadherin^EC1-5^ compared to controls (Figure 2B, D, F, H).

**Figure 2:**
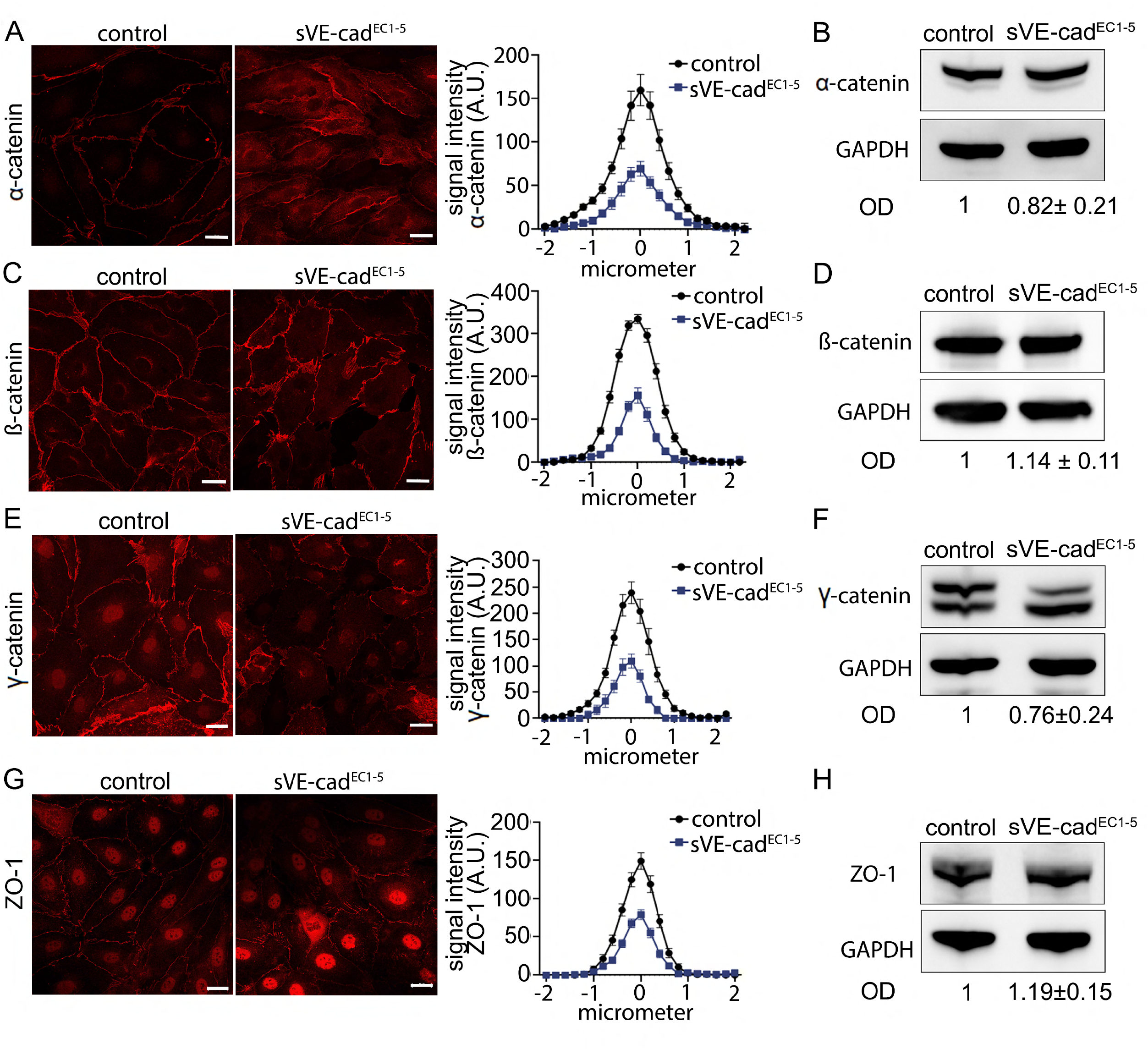
Cell junction proteins associated with VE-cadherin and tight-junction associated protein ZO-1 are reduced at the cell border in response to sVE-cad^EC1-5^. Immunofluorescence staining for a-catenin (A), b-catenin (C), g-catenin (E) and ZO-1 (G) of endothelial monolayer under control conditions and after 24h treatment with sVE-Cad^EC1-5^; changes were confirmed by quantification of the staining intensity (p<0.0001; n>5 for each of the conditions); scale bar is 20 μm. Representative Western Blot and quantitative analysis of whole protein levels of a-catenin (B), b-catenin (D), g-catenin (F) and ZO-1 (H) showed no differences.

### sVE-cadherin^EC1-5^-induced loss of endothelial barrier function is caused by loss of adhesion and VE-PTP-mediated signaling

RNA sequencing analyses following application of sVE-cadherin^EC1-5^ revealed minor changes of RNA expression levels for changes CDH5 (VE-cadherin), and TJP1 (ZO1) compared to controls (adj. p= 0.160 for CDH5 and adj. p=0.31 for TJP1). RNA expression levels of PTPRB (VE-PTP) were reduced (adj. p-value 0.00025 for PTPRB)(Figure 3A).

**Figure 3:**
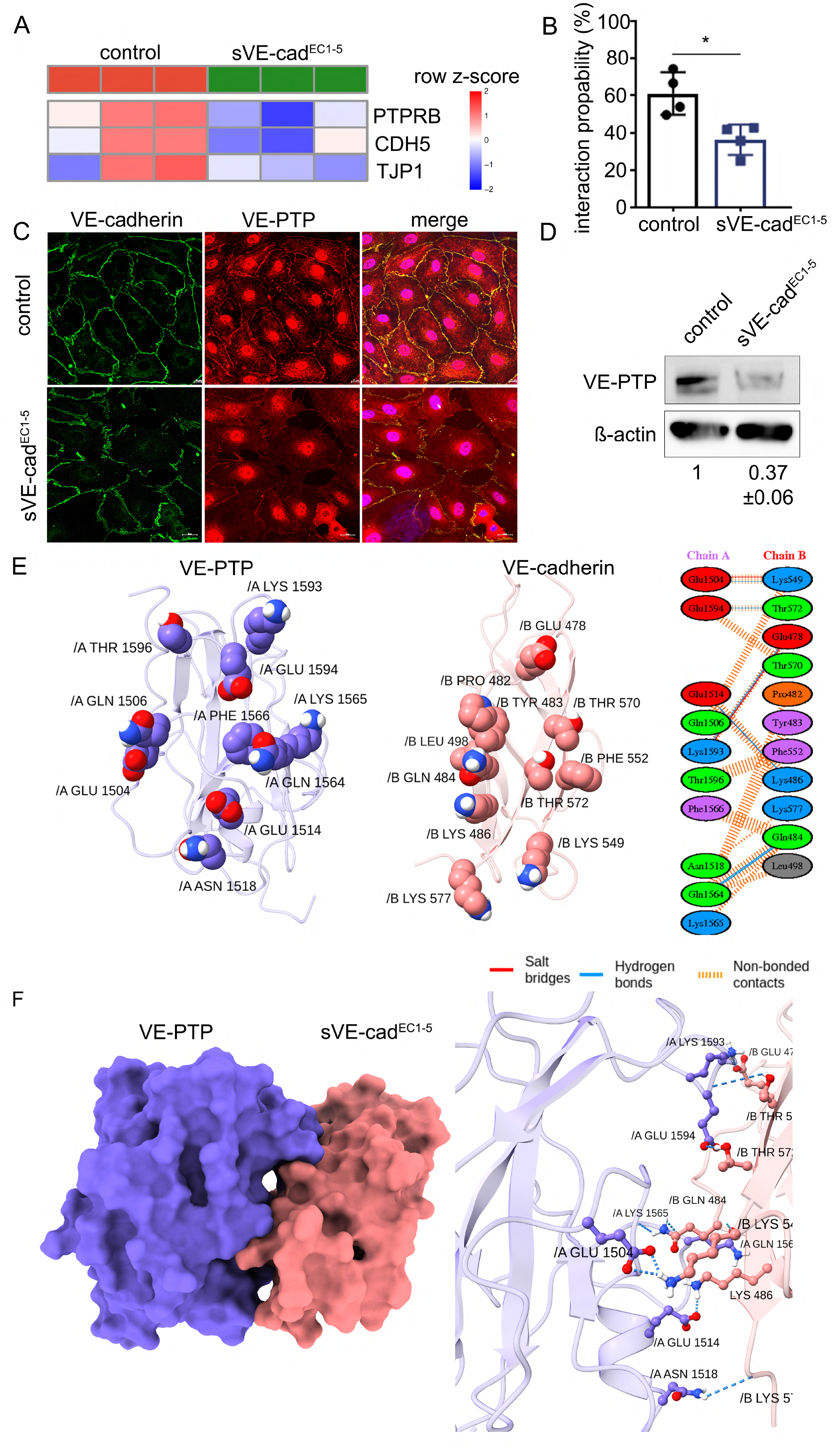
Interaction between VE-PTP and VE-cad^EC1-5^. A) Heatmap analysis summarizing the results of RNA-sequencing analyses for PTPRB (VE-PTP), CDH5 (VE-cadherin), and TJP1 (ZO1); n= 3 B) Atomic force microscopy experiments demonstrated that application of sVE-cad^EC1-5^ resulted in significantly reduced binding activity of VE-cadherin compared to control (p<0.012, n=4, paired two-tailed t-test). C) Co-immunostaining of VE-cadherin and VE-PTP showed a colocalization at the cell border under control conditions which was overall reduced following sVE-cad^EC1-5^; n=6 D) Western blot analyses of HDMEC lysates incubated with sVE-cad^EC1-5^ showed reduced VE-PTP protein levels compared to untreated cells (p<0.0001, n=5, unpaired two-tailed t-test). E) and F) Molecular dynamic simulations revealed an interaction between the 5^th^ EC-domain of VE-cadherin with VE-PTP.

To test whether the presence of sVE-cadherin^EC1-5^ interferes with the homophilic VE-cadherin transinteraction we used atomic force microscopy (AFM) where recombinant VE-cadherin molecules were covalently coupled to the cantilever tip and a mica sheet which were repeatedly brought into contact to allow assertion of the binding probability (Heupel et al., 2009). Application of sVE-cadherin^EC1-5^ resulted in a reduction of binding activity from 61±5% to 36±4% (Figure 3B).

Co-immunostaining of VE-cadherin and VE-PTP in endothelial cells showed a regular distribution at the cell periphery under control conditions. Application of sVE-cadherin^EC1-5^ to endothelial monolayers resulted in a fragmented staining pattern of both proteins (Figure 3C). Reduced co-localization suggested that the loss of interaction between VE-cadherin and VE-PTP could be a critical mechanism for endothelial barrier breakdown in response to sVE-cadherin^EC1-5^. According to RNA sequencing analysis, VE-PTP protein levels were reduced in Western blot analysis (following incubation with sVE-cadherin^EC1-5^ (Figure 3D).

In view of these observations we hypothesized that sVE-cadherin^EC1-5^ may also bind VE-PTP to compete with binding of VE-cadherin on the cell surface which is a critical interaction to maintain endothelial barrier properties (Vestweber, 2021). Molecular dynamic simulations were used to test whether the 5^th^ EC-domain of sVE-cadherin^EC1-5^ would bind VE-PTP to substantiate this hypothesis. Following identification of active amino acids as revealed by the amount of a surface area exposed to water for more than 40%, docking simulations at defined conditions were carried out. According to this, the number of amino acids in the VE-PTP and sVE-cadherin^EC1-5^ interfaces is 10 and 11, respectively (Figure 3E). Three salt bridges are formed between the two proteins and there are 6 hydrogen interactions and 59 non-bonded contacts between the two proteins. When studying the amount of van der Waals and electrostatic energies between the two chains during the molecular dynamics simulation the calculation of the mean value showed that the values of van der Waals and electrostatic energies are on average equal to −202.3 and −421 kg/mol, respectively. In summary, the simulation data suggested that sVE-cadherin^EC1-5^ may bind VE-PTP (Figure 3E, F).

Given that the loss of interaction between endothelial VE-cadherin and VE-PTP would result in phosphatase activation we tested whether pharmacological inhibition of the catalytic activity of VE-PTP using AKB9778 (Razuprotafib) would attenuate the effects sVE-cadherin ^EC1-5^ (Shen et al., 2014). As revealed by immunostaining of endothelial monolayers, VE-cadherin staining patterns were restored, stress fiber formation was reduced and no intercellular gap formation was evident when sVE-cadherin^EC1-5^ and AKB9778 were applied together compared to incubation with sVE-cadherin^EC1-5^ incubation alone (Figure 4A). AKB9778 alone showed no significant changes when compared to controls. In measurements of TER across endothelial monolayers co-incubation of sVE-cadherin^EC1-5^ together with AKB9778 abrogated the barrier-compromising effects of sVE-cadherin^EC1-5^ (sVE-cadherin^EC1-5^ 0.44 ± 0.08-fold of baseline; sVE-cadherin^EC1-5^ + AKB9778 0.77 ± 0.08-fold of baseline after 11 hours) (Figure 4B). Application of AKB9778 alone resulted in increased TER. Previously, it was demonstrated that barrier-protective effects of AKB9778 were dependent on consecutive Tie2-activation (Frye et al., 2015). Therefore, we tested whether direct activation of Tie2 using Angiopoetin-1 would abolish the effects of sVE-cadherin^EC1-5^. However, as revealed by TER measurements 500 ng/ml of Angiopoetin1 did not change sVE-cadherin^EC1-5^-induced loss of TER in endothelial cells (Figure 4C). A possible explanation for this is that sequencing analyses of the Tie2 pathway showed that Tie2 (TEK) was downregulated in response to sVE-cadEC1-5 (adj. p-value 2.68×10^-8^). In addition, we observed minor changes in ANGPT2 (Angiopoetin2; adj. p-value 0.848) and ANGPT1 (Angiopoetin1; adj. p-value 0.099) expression (Figure 4D). This suggested that other pathways than VE-PTP-mediated Tie2 activation are critically involved downstream.

**Figure 4:**
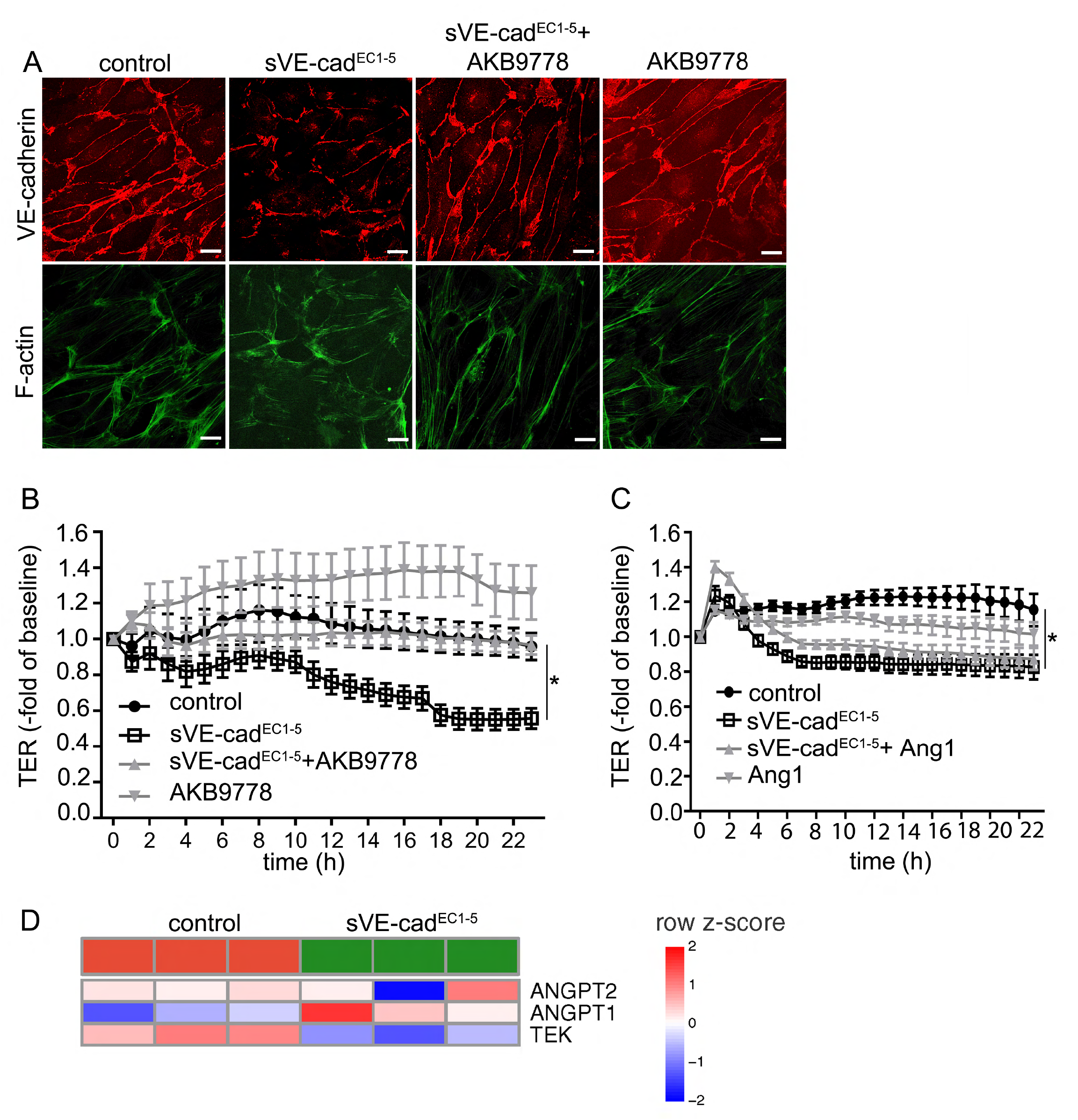
VE-PTP inhibitor (AKB9778) attenuated the effects of sVE-cad^EC1-5^. A) Immunofluorescence staining of VE-cadherin and F-actin in endothelial monolayers (HDMEC); images are representatives for n= 6, scale bar is 20 μm. B) and C) TER experiments using HDMECs under different conditions are shown (*p<0.0001, n=12, one-Way ANOVA with Friedman and Dunn’s Test). D) Heatmap analysis summarizing the results of RNA-sequencing analyses for Tie2 (TEK), ANGPT2 (Angiopoetin2) and ANGPT1 (Angiopoetin1).

### sVE-cadherin activates RhoA/ROCK pathway in VE-PTP-dependent manner

The strong formation of stress fibers in the actin cytoskeleton in response to sVE-cadherin^EC1-5^ led to the hypothesis that RhoA/Rho-kinase signaling may be involved downstream of VE-PTP-mediated signaling. Therefore, we used Rho-kinase inhibitor Y27632 in combination with sVE-cadherin^EC1-5^ in TER measurements across endothelial monolayers. These experiments demonstrated that Y27632 attenuated the effects of sVE-cadherin^EC1-5^ on endothelial barrier function (sVE-cadherin 0.5 ± 0.01-fold of baseline; sVE-cadherin+Y27632 1.07 ± 0.01-fold of baseline after 24 hours; Figure 5A). Application of Y27632 alone did not alter TER compared to controls (Y37632 1.07 ± 0.02-fold of baseline). In immunostaining the sVE-cadherin^EC1-5^-induced loss of VE-cadherin at cell borders was blocked by combined treatment of sVE-cadherin^EC1-5^ together with Y27632. Stress fiber formation and intercellular gap formation observed after sVE-cadherin^EC1-5^ alone were not observed following combined treatment of Y27632 and sVE-cadherin^EC1-5^ (Figure 5B). In Western blots for GEF-H1 we observed an overall increased expression following incubation of endothelial cells with sVE-cadherin^EC1-5^ (sVE-cadherin 1.43 ± 0.067-fold of control) (Figure 5C). GEF-H1 is known to promote the exchange of RhoA GDP to GTP and serves as a marker for RhoA activity. Next, we tested whether RhoA activity was increased in response to sVE-cadherin^EC1-5^ using pulldown assays (Figure 5D). As seen in Western Blot analyses of GEF-H1 protein expression after pulldown assay, application of sVE-cadherin^EC1-5^ increased GEF-H1 expression compared to untreated cells (2.7 ± 0.4 -fold of control). To confirm that RhoA/kinase signaling occurs downstream of VE-PTP-induced signaling we used AKB9778 together with sVE-cadherin^EC1-5^ Administration of AKB9778 blocked sVE-cadherin^EC1-5^-induced RhoA activation (1.2 ± 0.4 -fold of control) (Figure 5D, E).

**Figure 5:**
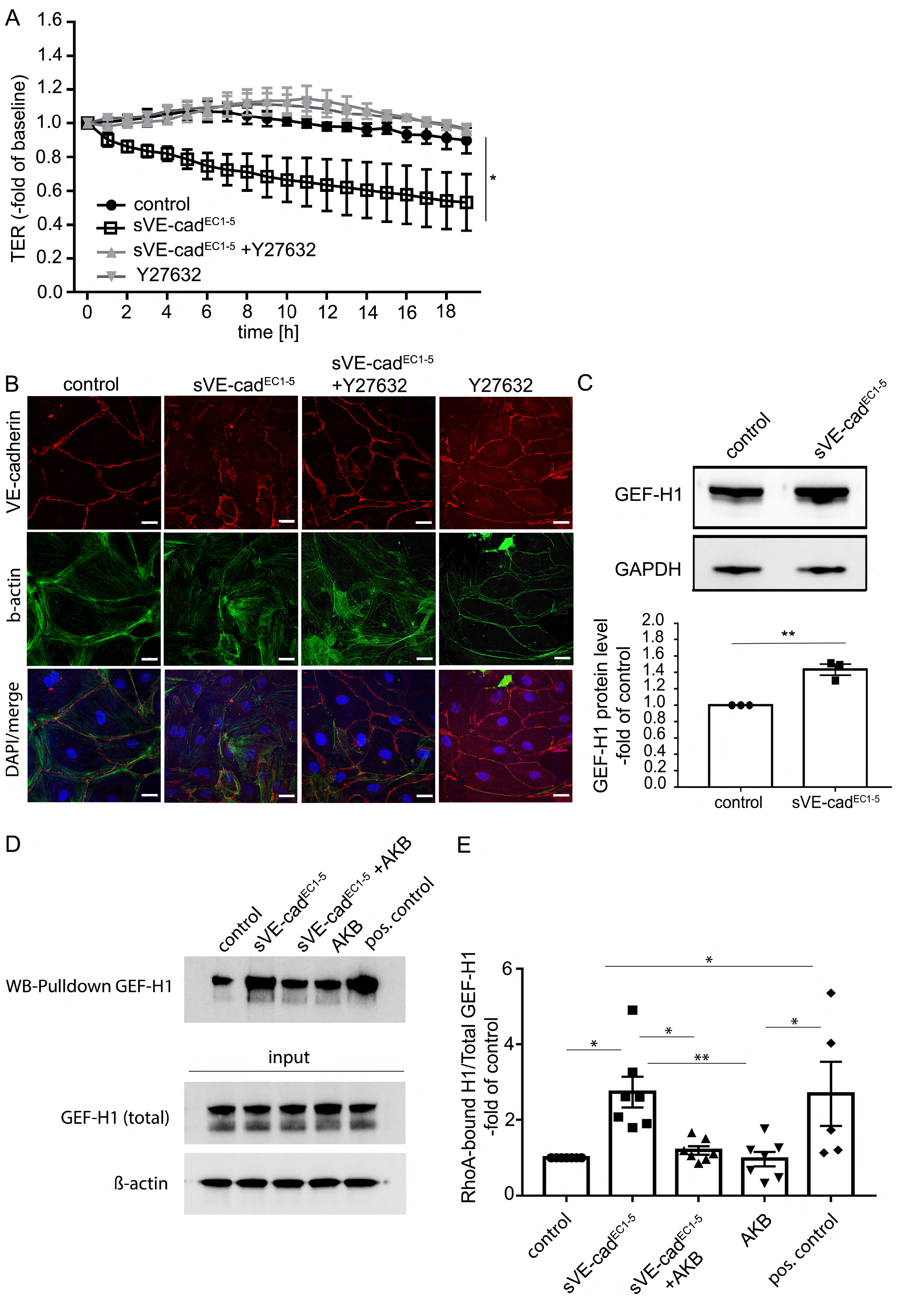
sVE-cad^EC1-5^ effects are mediated by VE-PTP-dependent activation of RhoA/ROCK pathway. A) TER measurements using HDMECs are shown (* p<0.001, n=4, ordinary 1-way ANOVA). B) Immunostaining of endothelial monolayers (HDMECs) for VE-cadherin (red) and F-actin (green) are shown; n=5; scale bar is 20 μm. C) Representative Western Blot and quantitative analyses of HDMEC whole cell lysates displayed increased GEF-H1 protein levels (marker for activated RhoA) after 24h incubation with sVE-cad^EC1-5^ compared to control conditions (p<0.0029, n=3, unpaired two-tailed t-test) D) Western Blots following pulldown of activated RhoA revealed an increased GEF-H1 protein level after application incubation with sVE-cad^EC1-5^ (p<0.01, n=7, ordinary 1-way ANOVA) compared to untreated cells. Application of AKB9778 blocked RhoA activation (p<0.01, n=7, ordinary 1-way ANOVA) whereas AKB9778 alone had no effect. As a loading control GEF-H1 Western Blot of whole cell lysate was performed before pulldown assay (GEF-H1 total) in untreated cells.

### sVE-cadherin disrupts microvascular barrier function and microcirculatory flow in vivo

To test the relevance of our observations *in vivo* we used a rat model that we had previously established to simultaneously monitor macro-hemodynamic and microvascular changes in systemic inflammation (Schick et al., 2012). In this model system we applied sVE-cadherin^EC1-5^ intravenously without additional administration of cytokines. The sequence of EC1-5 of VE-cadherin is 86% homologue between human and rats. Simulation of the structure of EC1-5 (Figure 6A) and calculation of the mean root square deviation (MRSD) between human and rat VE-cadherin using Chimera software version 2021, with Needleman-Wunsch alignment algorithm and BLOSUM-62 Matrix resulted in a RMSD of 0.306 angstroms (Å). An MRSD of less than about 2Å is generally considered as very close so that this justified to use the human sVE-cadherin^EC1-5^ construct for the *in vivo* experiments in rats.

**Figure 6:**
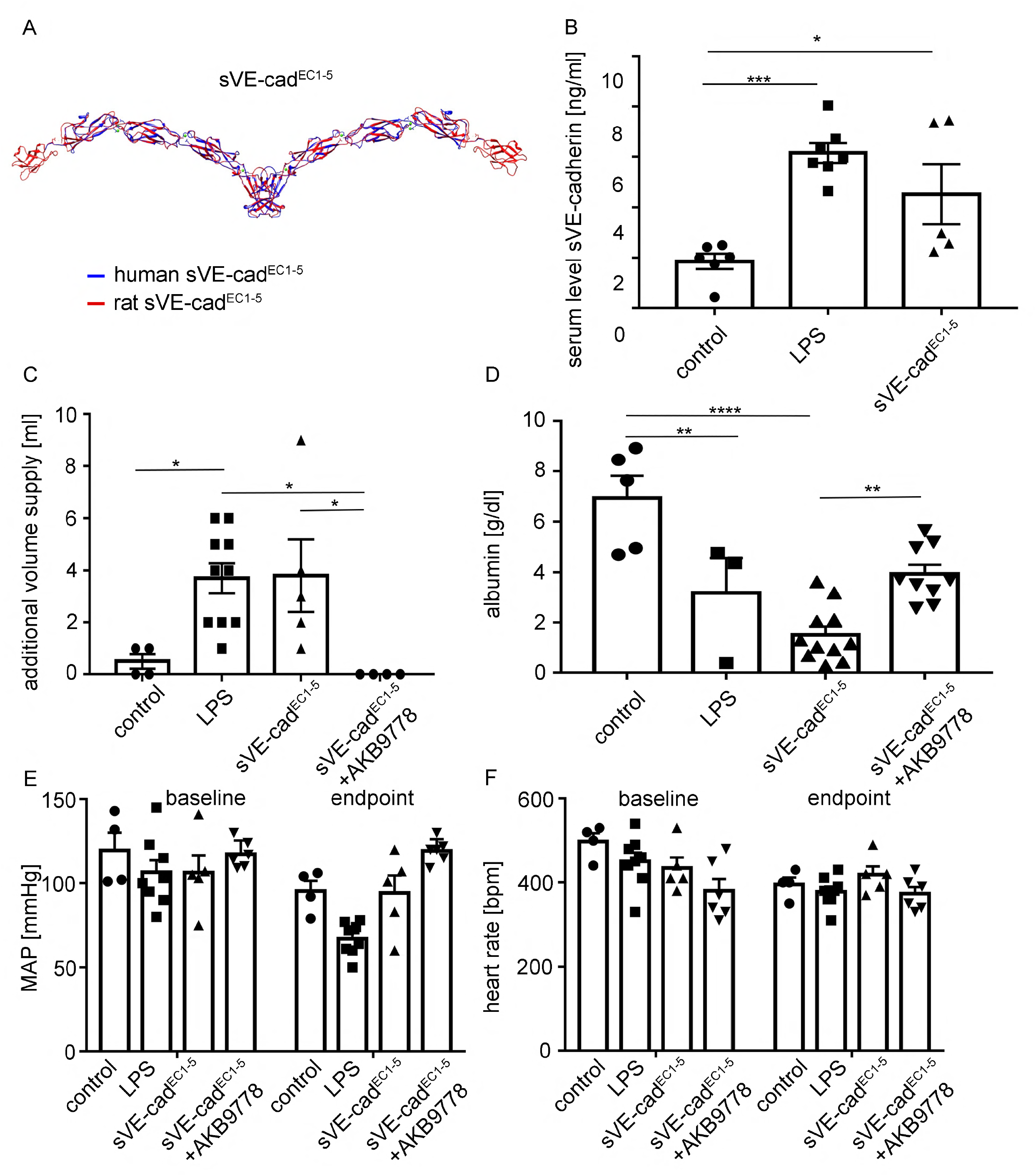
Application of sVE-cad^EC1-5^ altered macrohemodynamic parameters. A) Homology model of human and rat sVE-cad^EC1-5^ displayed, showing the highly similar structure resulting from their 86% sequence identity. B) Injection of either LPS or sVE-cadherin^EC1-5^ caused a significant increase in sVE-cad serum level when compared to control group (*p<0.005, ***p<0.001, n=5, ordinary 1-way ANOVA). C) Both LPS and sVE-cad^EC1-5^ treated rats required significantly more additional volume compared to control group (*p<0.05, n=5, ordinary 1-way ANOVA). D) Measurement of albumin in the serum at the end of the experiment revealed decreased albumin concentration in the LPS and sVE-cad^EC1-5^ group compared to control (*p<0.05, ****p<0.0001, n=5, ordinary 1-way ANOVA). E) and F) MAD and heart rate were documented throughout the whole experiment without differences.

In parallel, we used LPS in another group to induce systemic inflammation to compare this to animals subjected to sVE-cadherin^EC1-5^ alone. We assessed administration of sVE-cadherin^EC1-5^ by measurements of sVE-cadherin levels in the serum of the different groups after experiments were terminated. In control animals which received NaCl only, basal sVE-cadherin levels amounted 1.9 ± 0.9 ng/ml (Figure 6B). The i.v.-application of sVE-cadherin^EC1-5^ in the sVE-cadherin^EC1-5^ group resulted in augmented serum levels of sVE-cadherin of 4.5 ± 0.9 ng/ml. In animals following LPS application levels of sVE-cadherin were also elevated to 6.2 ± 0.9 ng/ml confirming the release of sVE-cadherin under conditions of systemic inflammation and that physiologically relevant doses had been applied in the sVE-cadherin^EC1-5^-treated animals.

Due to the experimental setup, which requested a fluid replacement with 1ml 0.9% NaCl intravenously when mean arterial pressure (MAP) dropped below 60 mmHg, MAP and heart rates were not different when controls were compared to the group of animals receiving sVE-cadherin^EC1-5^ or LPS before experiments were terminated. However, animals following sVE-cadherin^EC1-5^ application (3.8 ± 1.4 ml) and LPS-treated animals (3.7 ml ± 0.6 ml) required more additional i.v. volume application to maintain MAP above 60 mmHg than controls (0.5 ± 0.3 ml compared to control). In contrast, requirement of additional volume was not observed in the group of animals that received AKB9778 + sVE-cadherin^EC1-5^ (0 ± 0 ml compared to control) (Figure 6C). Serum albumin levels at the end of the experiments were 6.9 ± 0.9 g/dl in controls and were reduced to 1.5 ± 0.3 g/dl in sVE-cadherin^EC1-5^ group, 3.2 ± 1.4 g/dl in the LPS group, which suggested loss of microvascular barrier function and dilution due to volume replacement in the latter groups. This was not observed in animals treated with AKB9778 + sVE-cadherin^EC1-5^ (2.7 ± 0.7 g/dl compared to control) (Figure 6D).

No significant differences were seen in all treatment groups concerning MAP and heart rate at the beginning (baseline) and at the end of the experiment (Figure 6E, F). Measurements of alveolar septa in H&E-stained lungs of the different groups showed that animals following application of sVE-cadherin^EC1-5^ and LPS had significantly increased thickness of alveolar septa compared to the control group which amounted 5.38 ± 2.4 μm in controls, 21.03 ± 2.2 μm in the sVE-cadherin^EC1-5^ group and 12.67 ± 2.1 μm in the LPS group. In contrast, in animals treated with AKB9778 + sVE-cadherin^EC1-5^ no changes in alveolar thickness compared to controls was evident (9.8 ± 1.5 μm) (Figure 7A, B).

**Figure 7:**
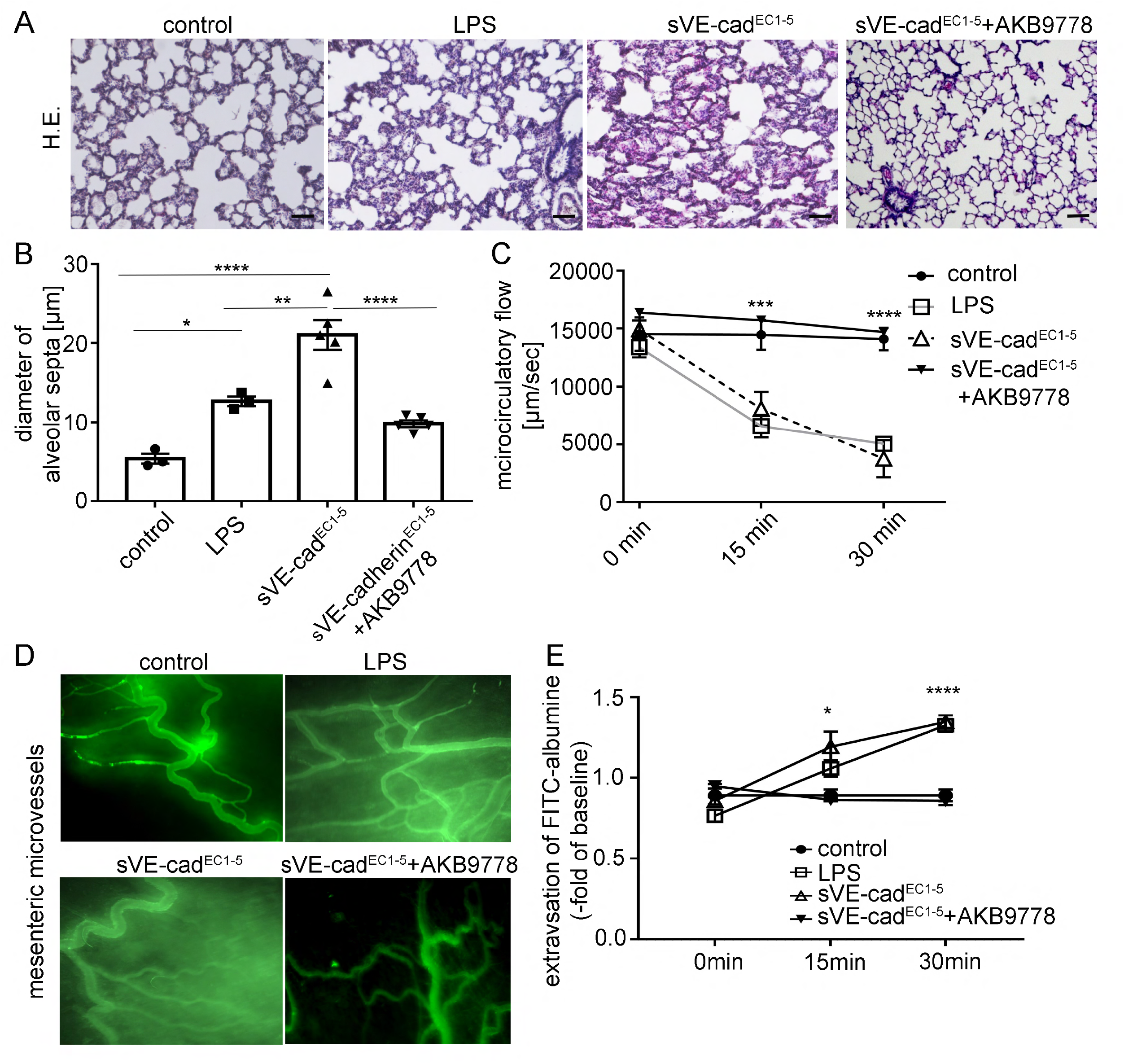
sVE-cadherin^EC1-5^ induces capillary leakage *in vivo*. A) Representative H&E-staining of lungs are shown. Scale bar is 100μm. B) Quantitative analysis of alveolar septa thickness is shown (*p<0.05, ***p<0.001, n=5, ordinary 1-way ANOVA). C) Microcirculatory flow was investigated by measurement of erythrocyte velocity [μm/sec] (n=5, ***p<0.001, ****p<0.0001, n=5). D) Representative pictures of FITC-albumin extravasation analyzed in E). E) FITC-albumin extravasation was quantified by measuring the change of light intensity (ΔI) inside and outside the vessels (*p<0.05, ****p<0.0001, n=5, ordinary 1-way ANOVA).

Microvascular blood flow as revealed by measurements of the velocity of erythrocytes within postcapillary mesenteric venules dropped significantly after 15min 0.56-fold of baseline (14460.3 ± 1281 μm/sec to 8087.3 ± 1447 μm/sec) in animals subjected to sVE-cadherin^EC1-5^ and to 0.45-fold of baseline (14460.3 ± 1281 μm/sec to 6588.8 ± 993 μm/sec) in animals subjected to LPS whereas microvascular flow remained unchanged in control animals over time (14519 ± 1434 μm/sec to 14090 ± 960 μm/sec) and in animals that were treated with AKB9778 + sVE-cadherin^EC1-5^ (14695 ± 875 μm/sec to 14460.3 ± 1281 μm/sec) (Figure 7C).

Investigation of microvascular capillary leakage was carried out by measurements of extravasation of i.v. injection of 5 mg/100 g body weight FITC-albumin at baseline, 15 and 30 min after the experiments were started. While no extravasation was observed in controls in the time course of the experiments and at the beginning of the experiments, application of sVE-cadherin^EC1-5^ resulted in increased extravasation of FITC-albumin to 1.3 ± 0.12 -fold of control after 15 min and to 1.5 ± 0.13-fold of control after 30 min. This was also the case in LPS-treated animals where extravasation of FITC-albumin increased to 1.2 ± 0.05 fold of control after 15 min and to 1.5 ± 0.06-fold of control after 30 min. In the group of animals receiving AKB9778 + sVE-cadherin^EC1-5^ no changes in FITC extravasation were observed (0.03 ± 0.01 fold of control after 15 and 0.03 ± 0.01-fold of control after 30 min (Figure 7D, E).

## Discussion

Formation of sVE-cadherin in inflammation has previously been linked with a poor prognosis in sepsis patients (Flemming et al., 2015; Yu et al., 2019; Yu et al., 2021; Zhang et al., 2010). However, it remained unexplored whether sVE-cadherin is just a result of inflammation or whether it also contributes to the vicious circle of endothelial dysfunction, loss of microvascular flow and organ failure in sepsis. This was addressed in the present study, where we used recombinant sVE-cadherin to test for its specific contribution to loss of endothelial barrier function which is commonly observed in patients with organ failure in sepsis. We found that sVE-cadherin targeted the VE-cadherin complex and led to stress fiber formation, intercellular gap formation and barrier dysfunction. The relevance of these observations in vivo was confirmed in a rat model where intravenous administration of sVE-cadherin increased microvascular permeability and disrupted microcirculatory flow. Our data provide evidence that sVE-cadherin disrupts endothelial barrier function by targeting the VE-cadherin/VE-PTP complex since all sVE-cadherin-induced effects were blunted by pharmacological inhibition of VE-PTP using AKB9778. Downstream signaling involves VE-PTP-dependent RhoA/Rho kinase activation. This points to a novel pathophysiological mechanism by which sVE-cadherin contributes to endothelial barrier disruption in vitro and in vivo. In addition, these data strengthen previous observations of a critical role of VE-cadherin/VE-PTP in endothelial barrier regulation in health and disease.

### sVE-cadherin is directly involved in loss of endothelial barrier function

Since intact VE-cadherin is required to maintain intercellular adhesion, it is well established that this represents a prerequisite to stabilize tight junction integrity and thereby plays the critical role in maintaining endothelial barrier integrity in vitro and in vivo (Corada et al., 2002; Dejana and Vestweber, 2013). Fragments of VE-cadherin have been reported to occur as a result of pro-inflammatory stimuli in endothelial cells by shedding e.g. via ADAM10 activation (Flemming et al., 2015; Schulz et al., 2008). Since fragmentation of VE-cadherin results in loss of intercellular adhesion it is reasonable that this represents a critical hallmark to induce loss of endothelial barrier function in inflammation (Radeva and Waschke, 2018). The role of circulating sVE-cadherin fragments in this context remained largely unexplored. Previous data from our own group gave rise to the hypothesis that circulating sVE-cadherin fragments may further contribute and aggravate loss of endothelial barrier function (Flemming et al., 2015). This is of pathophysiological relevance since microvascular leakage contributes critically to loss of microcirculation and thereby induces organ failure in patients with severe inflammation (Ince et al., 2016). To specifically address the question of a potential involvement of sVE-cadherin fragments in endothelial barrier dysfunction we cloned a recombinant sVE-cadherin fragment consisting of the EC1-5 domains which has previously been shown to be released following shedding events (Flemming et al., 2015; Schulz et al., 2008; Yu et al., 2019). In addition to these fragments it has also been described that smaller fragments are also released in response to inflammatory stimuli (Schulz et al., 2008). We focused here on the EC1-5 fragments since they appear to represent the major fraction of VE-cadherin fragments and have been proven to be associated with a poor outcome in patients with inflammation (Flemming et al., 2015; Schulz et al., 2008). As revealed by sequencing, Western blot analysis and specific commercially available ELISA measurements we confirmed that our cloning strategy truly resulted in the synthesis of sVE-cadherin EC1-5 fragments in CHO cells transfected with the vector.

The novel observation here is that the application of sVE-cadherin alone results in loss of endothelial barrier function in a dose-dependent manner in vitro and in vivo. This was shown to occur to a comparable extent as when LPS was applied to endothelial cells. The fact that sVE-cadherin^EC1-5^ was applied in the absence of bacterial toxins and/or proinflammatory stimuli proves the specificity of sVE-cadherin-induced effects.

In general loss of VE-cadherin commonly occurs in response to proinflammatory stimuli leading to reduced endothelial barrier functions (Radeva and Waschke, 2018; Spindler et al., 2010). Application of sVE-cadherin^EC1-5^ on endothelial monolayers led to loss of VE-cadherin at the cell borders and reduced total protein levels of VE-cadherin. Similarly, well known interaction partners of VE-cadherin including the junction-associated proteins α-, β-, γ-catenin that are critically involved in the proper maintenance of endothelial barrier function and are known interaction partners of VE-cadherin (Lampugnani et al., 1995) (Guo et al., 2008) (Iyer et al., 2004) were robustly decreased at the cell border as revealed by immunostaining. This redistribution is consistent with the disruptive effects of sVE-cadherin on endothelial barrier function while we found that sVE-cadherin treatment did not affect overall levels of these proteins in endothelial cells.

### sVE-cadherin^EC1-5^ interferes with VE-cadherin/VE-PTP interaction

When considering the strong effects of sVE-cadherin^EC1-5^ on endothelial monolayers it could be assumed that the presence of sVE-cadherin induces cell death. This, however can be excluded since neither apoptosis nor reduced cell viability were observed in response to sVE-cadherin^EC1-5^ application. When focusing on the potential mechanism underlying sVE-cadherin-mediated loss of endothelial barrier function it is tempting to be content with the experiments showing that sVE-cadherin^E1-5^ directly interferes homophilic binding as revealed by atomic force microscopy measurements. However, the strong changes of VE-PTP immunostaining patterns at the cell borders following sVE-cadherin^EC1-5^ led us to test the contribution of changes in VE-PTP-dependent signaling. Since previous data point to a critical role of VE-PTP/VE-cadherin interaction for endothelial barrier stability we speculated whether sVE-cadherin^EC1-5^ may interfere VE-cadherin/VE-PTP interaction (Frye et al., 2015; Nawroth et al., 2002). In this context, it is important to note that the interaction between both proteins is known to occur between the extracellular domain EC5 of VE-cadherin and the 17th domain of PTP protein (Nawroth et al., 2002). The use of molecular simulations clearly confirmed that sVE-cadherin^EC1-5^ may bind to VE-PTP which strengthened the hypothesis that sVE-cadherin fragments may compete with cellular VE-cadherin proteins and thereby induce endothelial barrier disruption. This is supported by immunostaining showing that colocalization staining patterns of VE-cadherin and VE-PTP are lost following application of sVE-cadherin^EC1-5^ on endothelial monolayers. A well-documented consequence of the dissociation of VE-cadherin/VE-PTP complex is the activation of different signaling pathways (Nottebaum et al., 2008). In line with this our experiments using AKB9778 (Razuprotafib) which blocks the catalytic activity of VE-PTP (Shen et al., 2014) showed that all functional and structural effects of sVE-cadherin^EC1-5^ application in endothelial cells were attenuated. This was also observed in the *in vivo* experiments. In a previous study VE-cadherin gene ablation did not counteract the barrier stabilizing effect of AKB9778 which led to conclusion that VE-PTP stabilizes endothelial junctions by VE-cadherin-independent mechanisms i.e. via Tie-2 (Frye et al., 2015). In addition it was shown that antibodies against the extracellular part of VE-PTP led to a dissociation and phosphorylation of Tie-2 (Winderlich et al., 2009). Since a similar effect in response to sVE-cadherin is conceivable we addressed a potential involvement of Tie-2 signaling by stimulating endothelial cells with different doses of Tie-2 activator Angiopoetin1. This however was not sufficient to attenuate sVE-cadherin^EC1-5^-induced loss of endothelial barrier function as revealed by TER measurements indicating that Tie-2 signaling is not critically involved in sVE-cadherin^EC1-5^-mediated loss of endothelial barrier function.

### RhoA signaling is involved downstream of VE-PTP activation in response to sVE-cadherin

Since we observed strong stress fiber formation in response to sVE-cadherin^Ec1-5^ we tested whether RhoA signaling may be involved (Radeva and Waschke, 2018; Spindler et al., 2010). Using immunostaining and TER measurements clearly confirmed that Rho kinase inhibitor Y27632 blunted sVE-cadherin^EC1-5^-induced effects. In line with this RhoGEF-H1 was augmented following application of sVE-cadherin^EC1-5^ and the GTPase RhoA was activated as revealed by pull-down assays. Finally, the observation that RhoA activity was prevented by AKB9778 indicates that it acts downstream of VE-PTP signaling. Thus, our data show that sVE-cadherin^EC1-5^ disrupts endothelial barrier function at least in part through VE-PTP/RhoA signaling and provide strong rationale to test VE-PTP and/or RhoA inhibitors in order to treat the endothelial barrier dysfunction associated with sepsis and possibly also other conditions. This supports previous observations showing that VE-PTP is involved in the regulation of RhoA activation by inhibition of RhoGEFs may be activated following VE-PTP activation (Juettner et al., 2019).

### Clinical implications of sVE-cadherin-induced effects

Observational clinical studies have associated increased systemic sVE-cadherin levels with pathological conditions ranging from acute kidney injury (Yu et al., 2019), systemic vasculitis (Chen et al., 2014), chronic spontaneous urticarial (Chen et al., 2017) and systemic sepsis (Flemming et al., 2015). We extended these investigations by showing in an *in vivo* rat model that sVE-cadherin increases vascular permeability and disrupts microvascular flow to a comparable extent as LPS. These findings provide *in vivo* evidence of a pathogenic role of sVE-cadherin in regulating vascular permeability. This has direct implications for the development of novel treatments for sepsis and other inflammatory conditions associated with increased vascular permeability. It can be speculated that the therapeutic reduction of sVE-cadherin levels could reduce severity of microvascular leakage and organ failure. This could be achieved either by preventing the shedding of VE-cadherin or by extraction of sVE-cadherin from the blood of septic patients using dialysis. In addition, the observation that AKB9778 is effective in preventing the effects of sVE-cadherin *in vitro* and *in vivo* warrants further investigations whether this could be the long-awaited therapeutic agent to stabilize endothelial barrier dysfunction in septic patients.

## Material and Methods

### Test reagents and antibodies

The following primary antibodies were used for Western blot (WB) or immunofluorescent (IF) staining as indicated below: Antibody against VE-cadherin extracellular domain (WB 1:1000, IF 1:50, R&D Systems, MAB9381), antibody against VE-cadherin intracellular domain (WB 1:1000, Santa Cruz, sc-9989), ZO-1 (WB 1:500, IF 1:50, Millipore, AB2272), α-catenin (WB 1:500, IF 1:50, Thermofisher, 71-1200), β-catenin (WB 1:1000, IF 1:50, Invitrogen, 71-2700), γ-catenin (WB 1:500, IF 1:50, Cell Signalling, 2309), rb-polyclonal VE-PTP against c-terminus (0.5 μg/ml, kindly provided by Dietmar Vestweber, Münster, Germany), cleaved caspase-3 (Asp175) antibody (WB 1:600, Cell Signaling, CatNo 9661), caspase antibody (WB 1:1000, cell signalling, CatNo 9662S), GAPDH (WB 1:5000, Sigma-Aldrich, G9295), β-actin (WB 1:3000, Sigma-Aldrich, A3854).

The following horseradish peroxidase-labeled IgG secondary antibodies were used for Western blotting: gam pox (WB 1:3000, Dianova, 115-035-003), garb pox (WB 1:3000, Dianova, 111-035-003), dam Alexa Fluor 488 (1:200, Invitrogen, A-21201), darb Alexa Fluor 488 (1:200, Invitrogen, A-21206), gam cy3 (1:600, Dianova, 115-165-003), garb cy3 (1:600, Dianova, 111-165-003).

### Cloning and generation of recombinant sVE-cadherin

sVE-cadherin^EC1-5^ was secreted into the supernatant by CHO cells that were transfected with the pcDNA-DEST47 containing the sequence of VE-cadherin EC1-5. Cloning was performed according to manufacturer’s protocol (Gateway cloning system, Thermo Fisher Scientific, Waltham MA, USA). First, the coding sequence of EC1-5 of human VE-cadherin was amplified using cDNA from HUVEC cells as a template. The used primer pairs were designed according the Uniprot Reference for human VE-cadherin (extracellular domain) (P33151 (CADH5_HUMAN)). The Gateway Technology uses the lambda recombination system to facilitate transfer of heterologous DNA sequences (flanked by modified *att* sites) between vectors. Two recombination reactions constitute the basis of the Gateway Technology. The first BP reaction facilitates recombination of an *att*B substrate (*att*B-PCR product of VE-cadherin EC1-5) with an *attP* substrate (donor Vector: pENTR/SD-D-TOPO vector (Thermo Fisher Scientific, Waltham MA USA)) to create an *att*L-containing ENTRY clone. The *attB* PCR primer were designed according to manufacturer’s protocol (Gateway cloning system, Thermo Fisher Scientific, Waltham MA, USA). Forward primer contains Shine-Dalgarno and Kozak consensus sequence for protein expression in both *E.coli* and mammalian cells. (Forward primer: 5’ GGG GAC AAG TTT GTA CAA AAA AGC AGG CTT CGA AGG AGA TAG AAC CAT GAT GCA GAG GCT CAT GAT GCT C 3’; Reverse primer: 5’ GGG GAC CAC TTT GTA CAA GAA AGC TGG GTC GTC AAA CTG CCC ATA CTT GAC 3’).

For the BP Recombination reaction, 10 μl of the amplified *att*B-PCR product, 2 μl pDONOR vector, 5 μl BP clonase reaction buffer, and 4 μl BP clonase enzyme is combined in a 1.5 ml microcentrifuge tube. After an incubation of 1 h at 25°C, 2 μl Proteinase K is added and again incubated for 1 h at 37°C. The created ENTRY clone (pENTR/SD-D-TOPO vector with VE-cadherin EC1-5 inserted) was transformed in competent *E.coli* cells, antibiotica selected and analyzed via colony PCR. The positive transformants containing the sequence of human VE-cadherin EC1-5 were used for the second recombination reaction, the LR reaction. The LR reaction facilitates recombination of an *attL* substrate (the created ENTRY clone) with an *att*R substrate (destination vector pcDNA-DEST 47 (Thermo Fisher Scientific, Waltham MA, USA)) to create an *att*B-containing expression clone. The Gateway pcDNA-DEST 47 Vector is a destination vector for cloning and exression of GFP fusion proteins in mammalian cells. Genes (VE-cadherin EC 1-5) in the created ENTRY clone are transferred to the destination vector backbone by mixing the DNAs with the Gateway LR Clonase II enzyme mix (Thermo Fisher Scientific, Waltham MA, USA). In detail: for the LR Recombination reaction, 10 μl of the created ENTRY clone, 2 μl destination vector, 4 μl 5X LR clonase reaction buffer, and 4 μl LR clonase enzyme is combined in a 1.5 ml microcentrifuge tube. The resulting recombination reaction is then transformed into *E.coli* and the expression clone selected. After an incubation of 1 h at 25°C, 2 μl Proteinase K is added and again incubated for 1 h at 37°C. The created destination vector (pcDNA-DEST47 vector with VE-cadherin EC1-5 inserted) was transformed in competent *E.coli* cells, antibiotica selected and analyzed via colony PCR. The positive transformants were used for all the following experiments.

### Transfection of CHO cells

Transfection reagents were prepared as follows: Reaction mix A containing 600 μl Opti-MEM (Thermo Fisher Scientific, Waltham MA, USA; Ref.: 31985062) and 24 μl Lipofectamine LTX reagent (Thermo Fisher Scientific, Waltham MA, USA) and reaction mix B containing 700 μl Opti-MEM, 14 μl Plus reagent and 8 μg of the pcDNA-DEST47-EC1-5 plasmid were mixed and incubated for 5 minutes at room temperature. Thereafter, CHO cells (70 % confluent) at transfection day were incubated with 700 μl medium and 300 μl transfection mixture. CHO cells were obtained from ATCC (PTA 3765).

Three to four days after transfection the supernatant was taken and Halt proteinase inhibitor cocktail (Thermo Fisher Scientific, Waltham MA, USA; Ref.: 78438) was added. The supernatant was stored at 4°C until usage. The concentration of EC1-5 in the supernatants was measured using sVE-cadherin ELISA.

### Soluble VE-Cadherin Enzyme-linked immunosorbent assay (ELISA)

ELISA-based quantification of soluble VE-cadherin in cell culture supernatants (R&D Systems, Wiesbaden-Nordenstadt, Germany, DCADV0) and rat blood (MYBioSource, San Diego, USA MBS2511473) samples was performed by using VE-cadherin ELISA according to manufacturer’s instructions. As maintained by the company the antibody in this ELISA recognizes epitopes between Asp48 and Gln593 which correspond to the extracellular domain of VE-cadherin (Flemming et al., 2015).

### Human primary cell culture of endothelial cells

We used Human Dermal Microvascular Endothelial Cells (HDMEC; Promocell, Heidelberg, Germany) that were described previously to be a suitable model to study microvascular endothelial barrier regulation in vitro (Flemming et al., 2015; Schlegel et al., 2009). Cells were used from passages 1 to 7 and grown in endothelial growth medium containing supplement mix and were passaged using Detach kit-30 (both Promocell). Culture confluency was checked by daily microscopy.

### Cell Viability Assay

For measuring cell viability CellTiter-Glo^®^ 2.0 Assay (Promega, Madison; USA; CatNo G9241), a luminescence-based cell viability assay was used. The number of viable cells in culture is determined by quantification of the amount of ATP present. The experimental set-up was designed and executed according to the manufacturer’s protocol. In brief, cells were grown on opaque-walled multiwall plates, CellTiter-Glo^®^ 2.0 reagent was added, and luminescence signal was measured. The luminescence signal is proportional to the amount of ATP present; the amount of ATP is directly proportional to the number of viable cells present in culture. Data were normalized to the controls of each experiment (Flemming et al., 2015).

### Measurement of transendothelial electrical resistance (TER)

ECIS 1600R (Applied BioPhysics, Troy, NY, USA) was used to measure the transendothelial resistance (TER) of HDMEC monolayers to assess endothelial barrier integrity as described in detail previously (Baumer et al., 2008). Endothelial cells were grown to confluence on 8 well arrays (8W10E+; Applied BioPhysics, Troy, NY, USA) and were incubated with fresh medium in the presence or absence of sVE-cadherin^EC1-5^ as indicated below. Additionally, Y27632 and AKB9778 were added simultaneously, when combined with sVE-cadherin^EC1-5^ at the concentrations described above.

### Permeability measurements across endothelial monolayers

HDMEC were seeded on top of transwell filter chambers on 12-well plates (0.4 μm pore size; Falcon, Heidelberg, Germany). After reaching confluence, cells were incubated with fresh medium without phenol red (Promocell, Heidelberg, Germany) containing 10 mg/mL FITC-dextran (70 kDa). Paracellular flux was assessed by taking 100 μL aliquots from the outer chamber over 2 h of incubation. Fluorescence was measured using a Tecan Microplate Reader (MTX Lab systems, Bradenton, USA) with excitation and emission at 485 and 535 nm, respectively. For all experimental conditions, permeability coefficients (P_E_) were calculated by the following formula as described previously (Schlegel et al., 2008): P_E_ = [(ΔCA/Δt) × VA]/S × ΔCL, where P_E_ = diffusive permeability (cm/s), ΔCA = change of FITC-dextran concentration, Δt = change of time, VA = volume of the abluminal medium, S = surface area, and ΔCL = constant luminal concentration (Meir et al., 2021).

### Immunocytochemistry

Immunostaining was performed as described previously in detail (Schlegel et al., 2008). HDMECs were grown until confluence on cover slips. After incubation with mediators as indicated below fixation with 2% formaldehyde for 10 min was performed. Cells were permeabilized with PBS including 0.1% Triton X-100 (Carl Roth GmbH + Co. KG, Karlsruhe, Germany, 3051.2) for 5 min and blocked in PBS containing 3% bovine serum albumin and 1% normal goat serum (Sigma-Aldrich, Taufkirchen, Germany, A4503 & G0023) at room temperature. Primary antibodies were applied overnight at 4°C. Coverslips were washed four times in PBS and then incubated with secondary antibodies for 1 h at room temperature in present or absence of Alexa Fluor 488 phalloidin (visualization of F-actin 1:60, Invitrogen, A-12379). Coverslips were then mounted using Vectashield HardSet™ with DAPI (Vector Laboratories, Inc.; Burlingame, USA) onto slides for fluorescence microscopy and captured using ZEISS LSM 780 microscope (Carl Zeiss AG, Oberkochen, Germany) and ZEN 3.4 software. Quantification of the images was performed as described previously (Burkard et al., 2021).

### Western Blotting

Cells growing on 6-well plates were scrapped after adding SDS lysis buffer. Lysates in Laemmli buffer were separated by SDS-PAGE and transferred onto nitrocellulose membranes by wet blotting. Blocking was carried out in 5% low fat milk in Tris-buffered saline Tween-20 (TBST). The membranes were incubated with the appropriate primary antibodies overnight at 4°C. Horseradish peroxidase labelled secondary antibodies were incubated for 1 h at room temperature. Proteins were visualized by the enhanced chemiluminescence technique (ECL, Amersham, Munich, Germany) and developed using ChemiDoc™ Imaging System (BioRad, Feldkirchen, Germany).

### Rho activity assay

For measurements of active RhoGTPase in cells, RhoA G17 agarose beads (Abcam, Cambridge, UK) were used according to manufacturer’s recommendations. Confluent HDMEC monolayers were lysed and incubated with the agarose beads, incubated for 2h at 4°C. After centrifugation and washing with RIPA buffer, the beads were resuspended in SDS-PAGE sample buffer and boiled for 5 min. The precipitated Rho-GEF was then detected by western blotting using GEF-H1 antibody (abcam, Cambridge, UK, Cat No ab155785). As an internal loading control, untreated equivalent aliquots of confluent HDMEC cells were harvested and western blot using GEF-H1 antibody was performed.

### Atomic force microscopy (AFM) measurements with recombinant VE-cadherin-Fc

AFM measurements were conducted on an atomic force microscope (Nanovizard III, JPK Instruments, Berlin, Germany) mounted on an inverted microscope (IX83, Olympus, Tokyo, Japan). AFM cantilevers (MLCT, Bruker, Calle Tecate, USA) and mica sheets (SPI supplies, West Chester, USA) were functionalized with recombinant VE-cadherin-Fc as outlined earlier (Heupel et al., 2009). Force-distance curves (FDC) were recorded with an applied force of 0.3 nN, a contact delay of 0.1 s and a retraction velocity of 1 μm/s. For each individual experiment, 1000 FDC were recorded in HBSS to assert baseline VE-cadherin trans-interaction. Thereafter, the mica and cantilever were incubated for 1h with 130 ng/ml sVE-Cad^EC1-5^ and 1000 additional FDC were recorded on the same position on the mica sheet. JPKSPM Data Processing software (JPK Instruments) was used to analyze recorded FDC and to determine VE-cadherin binding events. The resulting binding probability is the ratio of FDC that showed a VE-cadherin interaction and the total number of FDC.

### RNA extraction and sequencing

RNA from HDMEC was isolated using TRIZOL. RNA quality was checked using a 2100 Bioanalyzer with the RNA 6000 Nano kit (Agilent Technologies). The RIN for all samples was ≥7.7. DNA libraries suitable for sequencing were prepared from 300 ng of total RNA for four samples and appx 200 ng for two samples with lower concentrations, with oligo-dT capture beads for poly-A-mRNA enrichment using the TruSeq Stranded mRNA Library Preparation Kit (Illumina) according to manufacturer’s instructions (½ volume). After 15 cycles of PCR amplification, the size distribution of the barcoded DNA libraries was estimated ~325 bp by electrophoresis on Agilent DNA 1000 Bioanalyzer microfluidic chips. Sequencing of pooled libraries, spiked with 1% PhiX control library, was performed at 25 million reads/sample in single-end mode with 75 nt read length on the NextSeq 500 platform (Illumina) using High output sequencing kit. Demultiplexed FASTQ files were generated with bcl2fastq2 v2.20.0.422 (Illumina).

To assure high sequence quality, Illumina reads were quality- and adapter-trimmed via Cutadapt (Martin, 2011) version 2.5 using a cutoff Phred score of 20 in NextSeq mode, and reads without any remaining bases were discarded (command line parameters: --nextseq-trim=20 -m 1 -a AGATCGGAAGAGCACACGTCTGAACTCCAGTCAC). Processed reads were subsequently mapped to the human genome (GRCh38.p19) using STAR v2.7.2b with default parameters (Dobin and Gingeras, 2015). Read counts on exon level summarized for each gene were generated using featureCounts v1.6.4 from the Subread package (Liao et al., 2014). Multi-mapping and multi-overlapping reads were counted strand-specific and reversely stranded with a fractional count for each alignment and overlapping feature (command line parameters: -s 2 -t exon -M -O –fraction). The count output was utilized to identify differentially expressed genes using DESeq2 (Love et al., 2014) version 1.24.0. Read counts were normalized by DESeq2 and fold-change shrinkage was applied by setting the parameter “betaPrior=TRUE”. Differential expression of genes was assumed at an adjusted p-value (padj) after Benjamini-Hochberg correction < 0.05 and ļlog2FoldChangeļ ≤ 1. The data discussed in this publication have been deposited in NCBI’s Gene Expression Omnibus (Edgar et al., 2002) and are accessible through GEO Series accession number GSE210502: https://www.ncbi.nlm.nih.gov/geo/query/acc.cgi?acc=GSE210502.

### Experimental set-up of in vivo experiments

Experiments were performed on Sprague-Dawley rats (n=20; aged 5-7 weeks, 200-250 g; Janvier, 53940 Le Genest Saint Isle, France). Animals were kept under conditions that conformed to the National Institutes of Health “Guide for the Care and Use of Laboratory Animals”. All rats were maintained on a standard diet and water ad libitium for 12 h day and night cycles. Animals were not fasted prior to the procedure.

We combined established experimental set-ups to allow continuous measurements of microcirculatory flow and capillary leakage using intravital microscopy as well as the simultaneous assessment of macrohaemodynamic parameters to test the effects of sVE-cadherin in vivo (Schick et al., 2012; Sielenkämper et al., 2001; Waschke et al., 2004; Wunder et al., 2002).

Prior to 0.8-1.5% (v/v) isofluorane anesthesia (Forene Abbott, Wiesbaden Germany), 0.2 mg/kg BW butorphanol was injected subcutanously. To apply i.v. medication, the right jugular vein was cannulated. Additionally, the left carotid artery was cannulated for continuous blood pressure and heart rate measurements (Hewlett-PackardModel 88S, Hamburg, Germany). Tracheotomy was performed and rats were mechanically ventilated with FiO2 0.28 using a rodent ventilator (Type: 7025, Hugo Sachs Elektronic KG, March-Hugstetten, Germany) in a constant ventilation regime. Animals’ body temperature was always kept at 37°C using a heating plate. Sufficient depth of anaesthesia under the latter conditions had been tested in preliminary experiments and by estimation of heart rates, mean arterial pressure (MAP), leg movement in response to painful stimuli (which was possible because muscle relaxants were not applied) and eye-lid closing reflex.

Thereafter, median laparotomy was performed, and the mesentery was gently taken out and spread over a pillar as has been described elsewhere (Schick et al., 2012). The whole experimental set-up was then carefully placed under a modified inverted Zeiss microscope (Axiovert 200, Carl Zeiss, Göttingen, Germany) equipped with different lenses (Achroplan ×10 NA 0.25/×20 NA0.4/×40 NA 0.6). This allowed continuous observation of microcirculatory flow within postcapillary venules in the mesenteric windows as well as observation of capillary leakage following i.v. injection of FITC-albumin at 5 mg/100 g BW as described below. Images and videos were captured using a digital camera, ColorSnap CF (Photometrics, Tucson, USA), driven by Metamorph analysis software (Molecular Devices GmbH, Biberach a.d. Riss, Germany) and digitally recorded for off-line analysis as described below. In the time course of experiments the upper surface of the mesentery was continuously superperfused with 37.5°C crystalloid solution (Sterofundin, B. Braun Melsung AG, Melsung, Germany).

### Randomization of animals into experimental groups

After the preparation procedure animals were allowed to recover for 30 min and then samples for the first blood gas analyses (baseline samples) were taken using an ABL505 blood gas analyser (Radiometer, Copenhagen, Denmark). Each collected amount of blood was replaced by an equal volume of 0.9% sodium chloride (Fresenius Kabi, Bad Homburg, Germany). Before randomization of animals into different groups, the following criteria for exclusion were applied: MAP<60 mmHg, blood loss with Hb<7 g/dl, pH <6.9 or heart rate >550 bpm during preparation procedures. After baseline analyses of blood gas samples and haemodynamic parameters, rats were randomized into experimental groups. One group was used as control group (n=5), in one group (n=5) 130ng/ml sVE-cadherin^EC1-5^ adapted to the calculated blood volume/BW administered i.v. and one group received 130 ng/ml sVE-cadherin^EC1-5^ 1 h preincubated with VE-PTP inhibitor AKB 9778 (n=5) 0.6 mg per injection subcutaneously as described previously (Frye et al., 2015). An additional control group received 2.5 mg/kg BW LPS from *Escherichia coli* (Sigma, L2630; Deisenhofen, Germany) (n=10) to induce systemic inflammation (Schick et al., 2012).

### Intravital measurement of microvascular permeability

To assess changes of microvascular permeability during the experimental procedures, digital fluorescence images were taken after single i.v. injection of FITC-albumin 5 mg/100 g BW (Sigma, Deisenhofen, Germany). Fluorescence images were taken using a 100 W mercury lamp and a filter set consisting of a 450–490 nm excitation and a 520 nm emission filter within an inverted microscope (Zeiss Axiovert 200, Carl Zeiss, Göttingen, Germany). Images were taken before randomization, 15 min and 30 min (endpoint) after injection of either NaCl, sVE-cadherin^EC1-5^ or LPS or sVE-cadherin^EC1-5^ + AKB9778. Microvascular permeability was then estimated by determining the extravasation of FITC-albumin by measurements of integrated optical intensity as described previously (Bekker et al., 1989). The labelled FITC-albumin represented relative changes in permeability: *ΔI* =1-(*I*i Δ*I*o)/*I*i, where *ΔI* is the change in light intensity, *I*i is the light intensity inside the vessel, and *I*o is the light intensity outside the vessel. Grey scale values were measured in the postcapillary venules and in the extravascular space around the venules per unit area throughout the experiments and at selected times using ImageJ software. For each time point 10 randomly selected intravascular and interstititial areas near the postcapillary venules were selected for the measurements by a blinded observer.

### Evaluation of microcirculatory flow

Analysis of microvascular blood flow was carried out in straight segments of venular microvessels (20–35 *μm* in diameter). Velocity of erythrocytes was measured by a blinded investigator using the software VisiView (Puchheim, Germany). For each time point the velocities of 27 randomly chosen erythrocytes within postcapillary venules were measured for each animal. For each time point six to nine vessels per animal were analysed.

### Histopathological analyses of lungs

Lungs were removed after experimental procedures for histopathological studies. The tissues were fixed in formaldehyde 3.5% (Otto Fischar, Germany) for more than 24 h. Tissues were then embedded in paraffin and subsequently, sections were stained with H&E for analyses of morphological alterations within the tissues as described previously (Schick et al., 2012). BZ-II Analyzer Software was used to measure thickness of alveolar septa. In each of the animals, three sections were analysed and in each section thickness of 15 randomly chosen alveolar septa were measured by a blinded observer.

### Modelling of sVE-cadherin interaction with VE-PTP

3D structures of VE-PTP and VE-Cadherin were extracted from Alphafold databank respectively with P23467 and P33151 cods. Easy version of HADDDOCK online server was used to assess the potential interaction between sVE-cadherin (5^th^ domain) and the 17^th^ domain of VE-PTP. Amino acids with a surface area which exposed to water were more than 40%, were considered as active amino acids. After molecular docking, the output data were divided and the classes were rated based on Z-score. After performing molecular docking and selecting the best complex, the complex was used to do molecular dynamics simulation. To simulate molecular dynamics, the protonation state of ionisable atoms such as aspartate, glutamate, lysine, arginine and histidine at pH7 was first determined using Propka software. Gromacs 2021.5 software was used to simulate molecular dynamics and amber99SB + ILDN force field was used to parameterize the protein. The protein molecule was then placed inside an octagonal box and the distance of the protein from the walls of the box is 0.8 nm. The TIP3P water model added to the box as the solvent. And 4 sodium ions were replaced with water molecules to neutralize the system. The energy of the system was then minimized in 50,000 steps. The steepest descend algorithm was used for minimization. The system temperature then equilibrated to 300 K on the NVT stage for 200 ppm. V-rescale algorithm was used to balance the temperature at this stage. Then, in the NPT stage, the system was stabilized on one Bar pressure with the Parrinello-Rahman algorithm for 200 ps. In the mentioned equilibration steps, heavy protein atoms with a force of 1000 kJ / m2 were kept constant (kJ / mol nm2). In the final stage (production), the simulation was continued under NPT for 100 ns. The time step in all stages of the simulation is 2 fs. The LINCS algorithm was also used to keep the length of covalent bonds (other than hydrogen bonds with heavy atoms) constant. In the production stage, the Nose-Hoover algorithm was used to keep the temperature constant.

### Statistical analysis

Statistical analysis was performed using Prism (GraphPad Software, La Jolla, CA, USA). Data are presented as means ± SEM. Statistical significance was assumed for p<0.05. Paired Student’s t-test was performed for two-sample group analysis after checking for a Gaussian distribution. Analysis of variance (ANOVA) followed by Tukey’s multiple comparisons test and Bonferroni correction was used for multiple sample groups. For non-parametric data, Kruskal–Wallis following Dunn’s post-test or Mann–Whitney U-test were used for significant differences. The tests applied for each of the different experiments are indicated in the figure legends.

### Ethical statement and study approval

All animal experiments were approved by the animal care committee (laboratory animal care and use committee of the district of Unterfranken, Germany) by the Regierung von Unterfranken (AZ 2-1241), Germany. All animal experiments performed conform to the guidelines from Directive 2010/64/EU of the European Parliament on the protection of animals used for scientific purposes. Human endothelial cells used in this study were obtained commercially (HDMEC; Promocell, Heidelberg, Germany). The preparation of the endothelial cells conforms to the Declaration of Helsinki.

## Data availability

All data generated or analyzed during this study are included in the manuscript and supporting files. The data discussed in this publication have been deposited in NCBI’s Gene Expression Omnibus (Edgar et al., 2002) and are accessible through GEO Series accession number GSE210502: https://www.ncbi.nlm.nih.gov/geo/query/acc.cgi?acc=GSE210502.

## Author contributions

All authors have made substantial contributions to conception or design and approved the submission.

JLK and NB: performed experiments, analyzed data, interpreted data, involved in the conception and design of the work. Wrote the initial draft of the article.

MD: performed experiments, analyzed data, interpreted data

TD: performed experiments, analyzed data, interpreted data

MS: performed experiments, analyzed data, interpreted data

MH: performed experiments, analyzed data, interpreted data

JW: performed experiments, analyzed data, interpreted data

SF: performed experiments, analyzed data, interpreted data, involved in the conception and design of the work. Wrote parts of the initial draft of the article, third party funding.

NS: performed experiments, analyzed data, interpreted data, involved in the conception and design of the work, supervised the whole work, third party funding, wrote and finalized the manuscript and all figures.

## Acknowledgments

rb-polyclonal VE-PTP antibody was kindly provided by D. Vestweber.

---

**Figure legend suppl. Figure 1:**
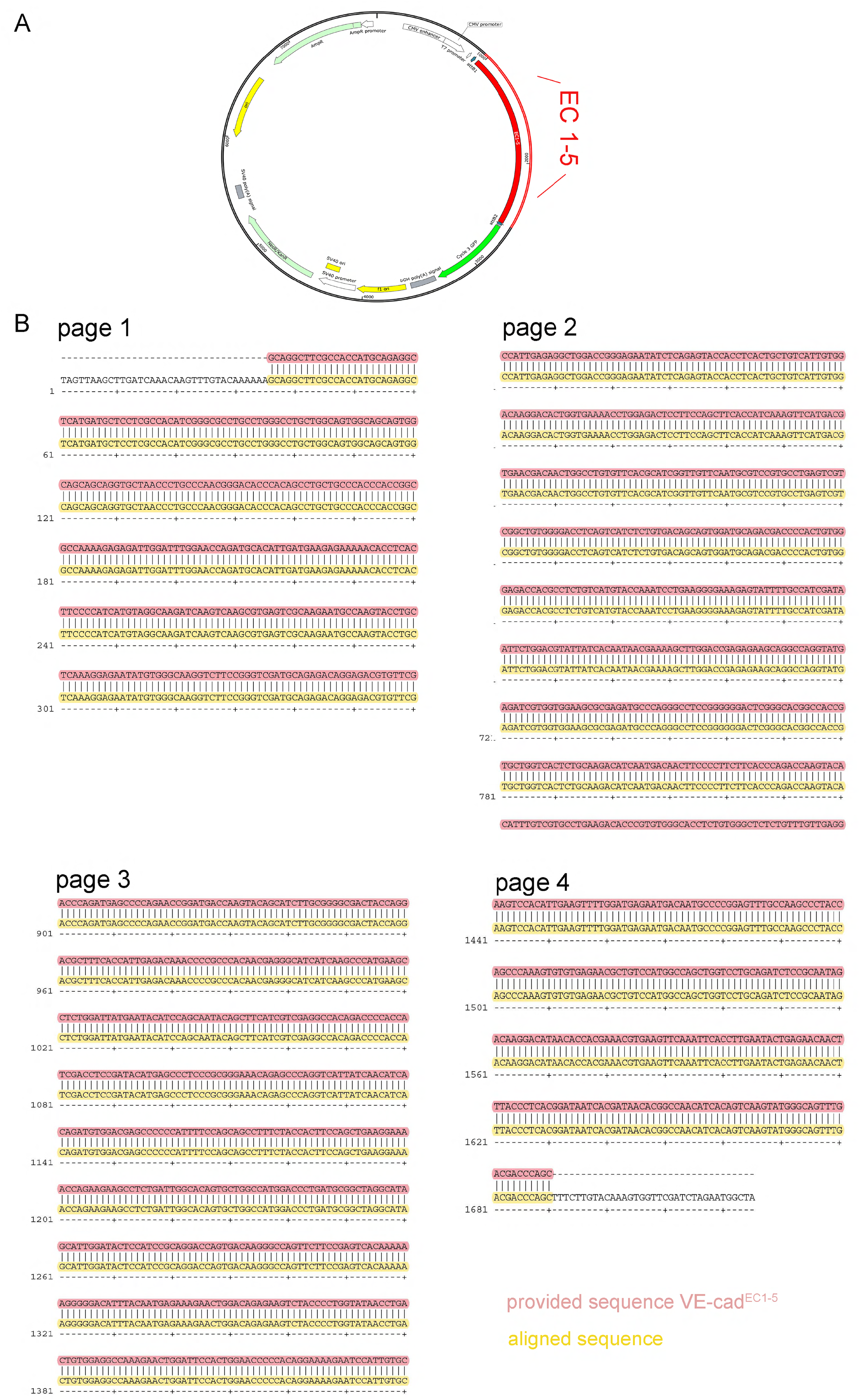
A) Schematic map of cloned pDEST47-VE-cadherin^EC1-5^ vector. The sequence of VE-cadherin EC1-5 was flanked via *attB-sites*. For antibiotic selection, the cloned vector contains resistance genes for kanamycin and neomycin. B) Alignment of cloned pDEST47-VE-cadherin^EC1-5^ displayed a sequence identity of 100%.

